# A Supervised, Symmetry-Driven, GUI Toolkit for Mouse Brain Stack Registration and Plane Assignment

**DOI:** 10.1101/781880

**Authors:** Marcelo Cicconet, Daniel R. Hochbaum

## Abstract

Immunostaining of brain slices is a ubiquitous technique used throughout neuroscience for the purposes of understanding the anatomical and molecular characteristics of brain circuits. Yet the variety of distortions introduced, and the manual nature of the preparation, hinder the use of the generated images from being rigorously quantified; instead most registration of brain slices is done laboriously by hand. Existing automated registration methods rarely make use of geometric shape information. When registering anterior-posterior brain slices, for example, small errors between consecutive planes accumulate, causing the symmetry axis of a plane to drift away from its starting position as depth increases. Furthermore, planes with imaging artifacts – e.g. one half of the slice is missing – can cause large errors, which are difficult to fix by changing global parameters. In this work we describe a method in which we register a set of consecutive brain slices enforcing all slices to have a vertical axis of symmetry, and then pair these slices optimally to planes from the Allen Mouse Brain Atlas via Dynamic Programming. The pipeline offers multiple human-in-the-loop opportunities, allowing users to fix algorithmic errors in various stages, including symmetry detection and pairwise assignment, via custom graphical interfaces. This pipeline enables large-scale analysis of brain slices, allowing this common technique to be used to generate quantitative datasets.

## 1 Introduction

This work describes – and serves as documentation for – an interactive toolkit aiming to register, plane-by-plane, a set of mouse brain slices to the Allen Mouse Brain Atlas^1^ [2], as well as quantifying puncta or difuse signal (e.g. from separate channels) in various labeled regions of the atlas.

The toolkit, available at github.com/hms-idac/RiffleShuffle, was developed using Matlab^2^ R2018a, and consists of a series of code cells, which the user executes sequentially, setting the appropriate parameters when needed, and adjusting possible algorithmic errors if necessary via custom designed graphical user interfaces.

While the pipeline can be executed with minimal interaction once the proper parameters are set, we found the ‘human in the loop’ approach to be critical due to the complexity of the task, specially when the amount of available data is small and therefore less forgetful to small errors – which in large datasets would possibly be dealt with by the law of large numbers.

Besides supervision, another feature that distinguishes this method from general registration algorithms is its assumption, and use of, bilateral symmetry. Indeed the anterior-posterior slices of the Allen Mouse Brain Atlas do display such property. Furthermore, assuming bilateral symmetry simplifies registration, since the dimension of the space of parameters is reduced. Finally, enforcing symmetry is a way to reduce the effect of error propagation in a sequence of pairwise registrations: in symmetry-agnostic methods, small errors between consecutive planes accumulate causing the symmetry axis of a plane to drift away from its starting position as depth increases.

There are three main phases: (1) pre-processing, (2) stack registration, (3) plane assignment. The pre-processing phase consists of resizing images, computing masks and edge maps, as well as detecting puncta if needed. Stack registration amounts to aligning the target dataset with itself using symmetry as a guide. In plane assignment the slices of the target dataset are paired to the slices of the atlas, starting from suggested *bregma* values, using a method similar to sequence alignment – based on dynamic programming; after pairing, target slices are further adjusted to Atlas slices using local non-linear deformations; finally, the amount of signal or spots intersecting each labeled brain area is quantified.

## 2 Methods

### 2.1 Pre-Processing

First, the data is downsized, primarily to reduce computational cost without affecting the subsequent steps. The pipeline assumes that each plane has two channels, one of which is used for registration, and the other for quantification. In principle, they could be the same. There are two options for quantification: puncta or diffuse signal. For diffuse signal, the user should keep in mind that no background subtraction, or flat-field correction, or any other image-acquisition controls are performed – they should be implemented beforehand if necessary. If the user chooses to quantify puncta instead, there are two code cells to inspect the approximate size of puncta, as well as testing how the chosen parameters for puncta detection work on selected image planes. Figure 1 has a sample output of the puncta-measurement interface.

**Figure 1:**
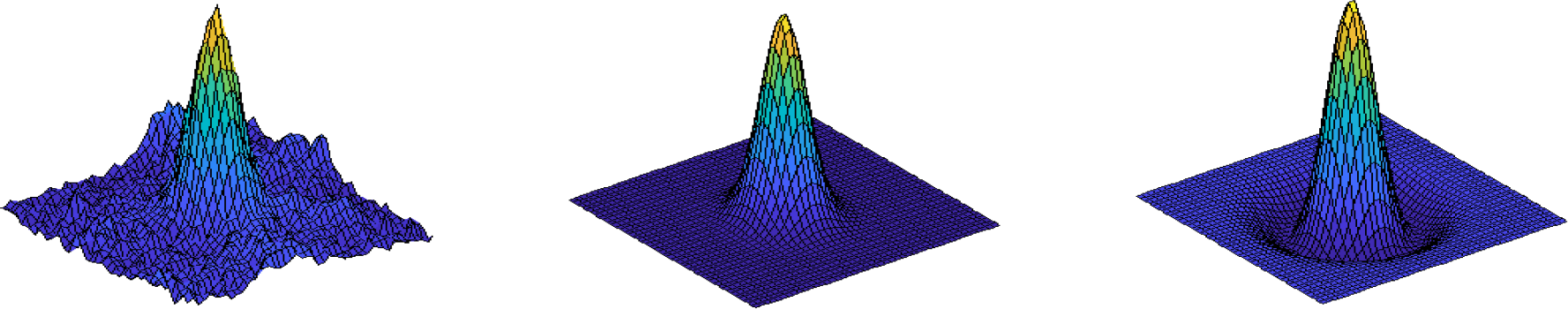
Output of the puncta-fitting GUI: pixel intensity map around puncta, reconstructed gaussian, and reconstructed laplacian of gaussian.

Measurement implies finding the sigma of the 2D gaussian that best-fits a rectangular crop around the puncta. For this we use a fitting algorithm developed by Nathan Orloff^3^.

Puncta detection is performed in two steps. First, a set of selected candidates is detected, based on finding local maxima in the output of a laplacian-of-gaussian filtering of the image. These candidates are then selected based on correlation to an ideal puncta (assumed to be a 2D gaussian with the estimated sigma). For the initial candidate selection, a mask that separates the brain slice from the background is used. This mask can be computed in various ways, but for the current implementation we trained a machine learning model to perform such segmentation.

At this stage we also compute ‘contour likelihoods’, which are heat maps that highlight the edges of the images. This is obtained using a separate machine learning model. These are the images we use for self-registration (step 2) and assignment (step 3). Using edge likelihoods is not only sufficient for registration, it is also helpful, since the target dataset typically has different fine details compared to the Atlas, which if taken into account would make non-linear registration difficult in step 3.

Figure 2 shows a sample output of this phase for a single plane.

**Figure 2:**
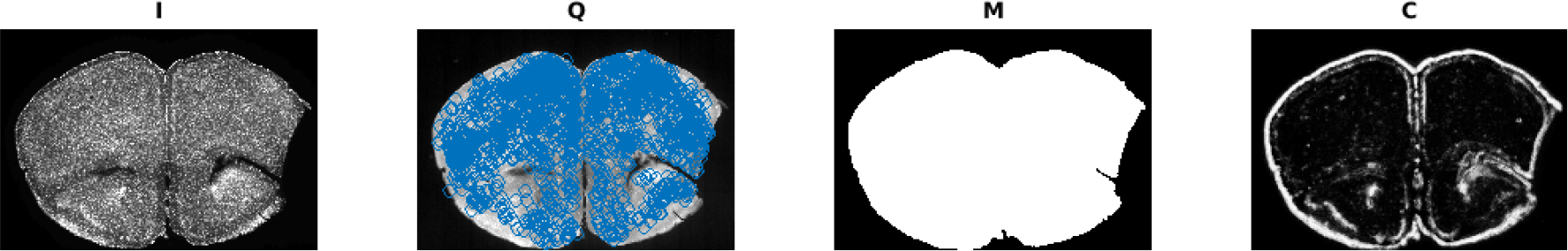
Pre-processing output. I: channel used to compute mask and contour likelihood; Q: channel to quantify, with locations of detected puncta overlayed; M: mask, C: contour likelihood.

### 2.2 Stack Registration

First, we run an automated symmetry detection algorithm [1] that finds the axis of bilateral symmetry in each plane independently and transforms the plane so that the new axis of symmetry coincides with the vertical line splitting the image in two equal halves. For planes where the algorithm fails to find the correct axis, adjustments can be made by running the global symmetry tool – Figure 3.

**Figure 3:**
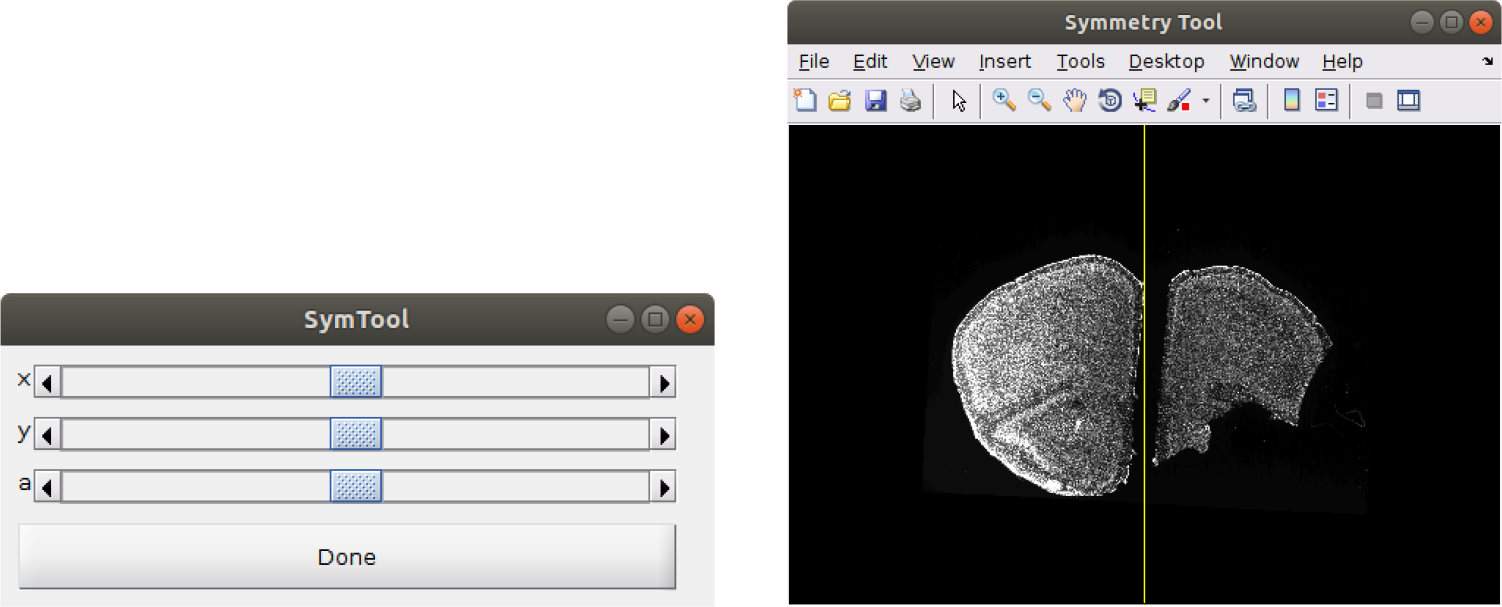
Controls and visualization for the global vertical symmetry adjustment tool.

In cases where a plane contains two partial brain slices, or even just one half of a slice, the user can further adjust each half independently using the partial symmetry tool – Figure 4.

**Figure 4:**
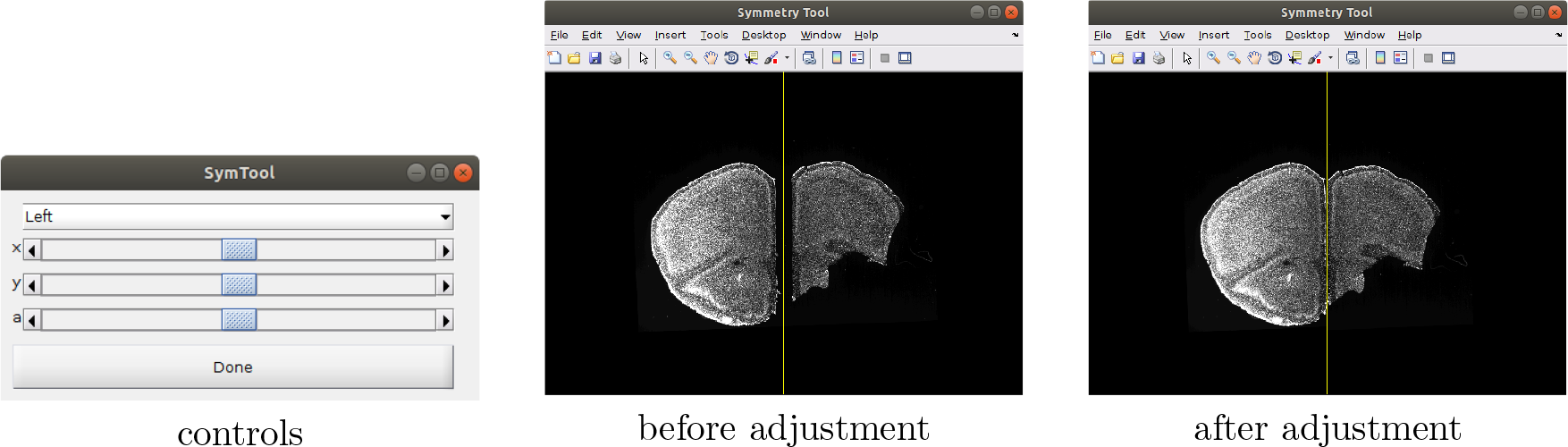
Controls and visualization for the partial vertical symmetry adjustment tool, where adjustments can be done independently to each half of the image.

This is followed by pairwise vertical registration, where each pair of consecutive planes is registered vertically. After this step the pairwise transformations are combined into a global vertical registration, where the middle plane is used as an anchor against which all planes are registered. If there is an error in pairwise vertical registration (leading to an error in global vertical registration), it can be fixed using a pairwise vertical registration tool – Figure 5.

**Figure 5:**
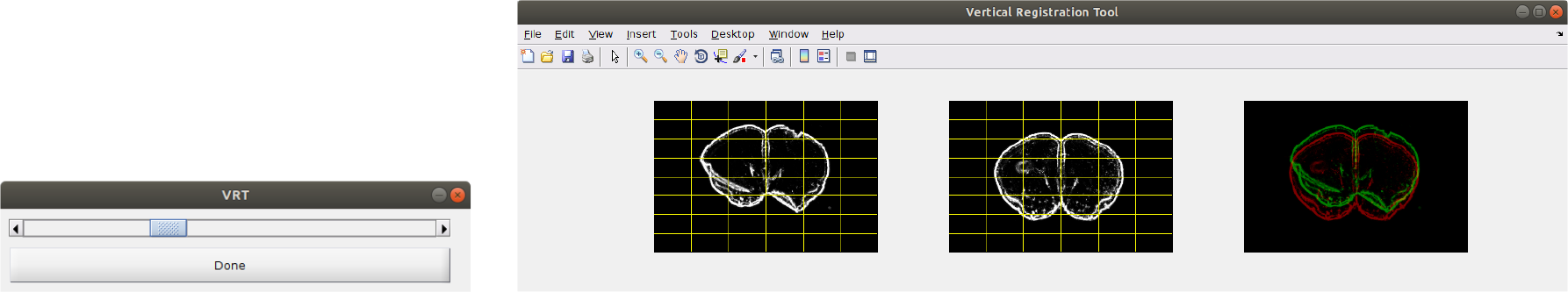
Pairwise vertical registration tool.

At this point, the dataset is *self-registered*, except for minor local deformations. During the steps above, corresponding masks and quantification channels (or puncta) are transformed accordingly. The results are saved to file for plane assignment to the Allen Mouse Brain Atlas.

### 2.3 Plane Assignment

We start by reading and resizing the Atlas, to approximately match the scale of the dataset we just worked on. Subsequently, the registered dataset (output of the previous phase) is loaded, and an appropriate region of the Atlas cropped in the *z* dimension. This is done based on the estimated range of bregmas covered by the dataset. For example, if the dataset contains slices with bregmas linearly spaced between *b*_0_ and *b*_1_, the Atlas is cropped so as to have planes only around those bregmas, with some buffer on each side – the amount controlled by an adjustable parameter. We further compute the 3D gradient magnitude of the Atlas, as this more closely resembles the edge likelihoods of the dataset. Figure 6 shows an example of the resulting planes from Atlas and dataset that become inputs to the pairwise assignment algorithm.

**Figure 6:**
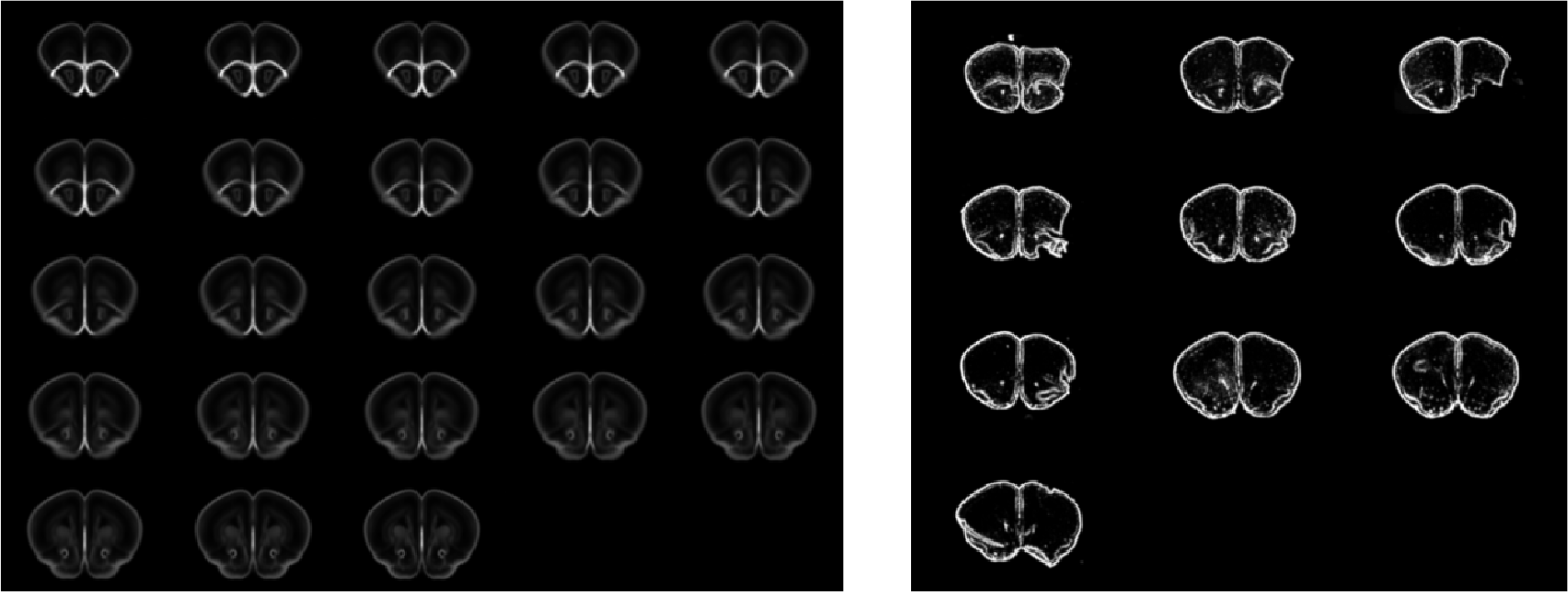
Left: planes of the subset of the Atlas to which the planes of the dataset will be aligned. Right: dataset planes after self-registration.

Before pairwise assignment, however, the user has the option to further resize the Atlas’ planes to match the scale of the dataset. Proper scaling increases both the accuracy and speed of the matching algorithm.

Plane assignment is performed using a dynamic programming algorithm that maximizes the sum of the plane-to-plane correlations. When computing candidates for pairing, the algorithm allows some further re-scaling and vertical displacement to measure the correlation between candidate pairs. Assignment is allowed within a certain range of the initially suggested positioning. For example, if the initial guess for the bregma value of a plane is *b*, the final assigned bregma should be within [*b* − *ϵ*, *b* + *ϵ*]. Figure 7 illustrates correlations for candidate matchings as well as final assignment.

**Figure 7:**
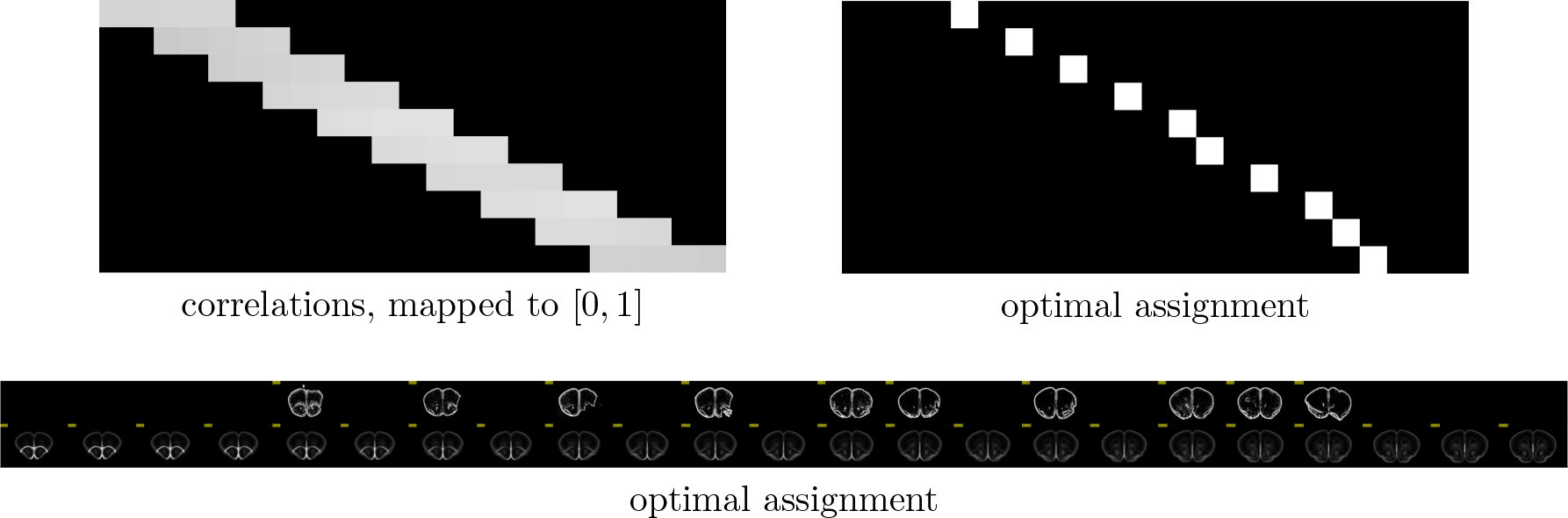
Outputs from the plane assignment algorithm. The top left image shows restrictions in bregma range. The top right image displays final assignment optimizing global correlation. Both represent Atlas planes horizontally and dataset planes vertically. The bottom image displays the actual plane assignment, as well as the possible candidates – this corresponds to the solution illustrated in the top right.

Results of the assignment are saved in a CSV table. If necessary, adjustments in this table can be done to fix assignment errors – by running the appropriate code cell in the Matlab script.

At this point, the datasets are largely aligned. However, due to manual nature of sample preparation, small deformations are introduced; thus an additional non-linear registration step is performed. Here we deploy a registration method based on Maxwell’s Demons [3, 4], as implemented by Matlab’s *imregdemons* function. Instead of using edge maps, however – as we did for the registration algorithms so far – we deploy *imregdemons* on binary images of the outer contours of the slices (see Figure 8). This helps the algorithm focus on small global adjustments, rather than errouneously stretch small areas to fit local features in images that originated from distinct imaging systems. Computation of the outer contours is based on steerable filtering and masks from the Atlas. Figure 8 illustrates inputs and outputs from this step.

**Figure 8:**
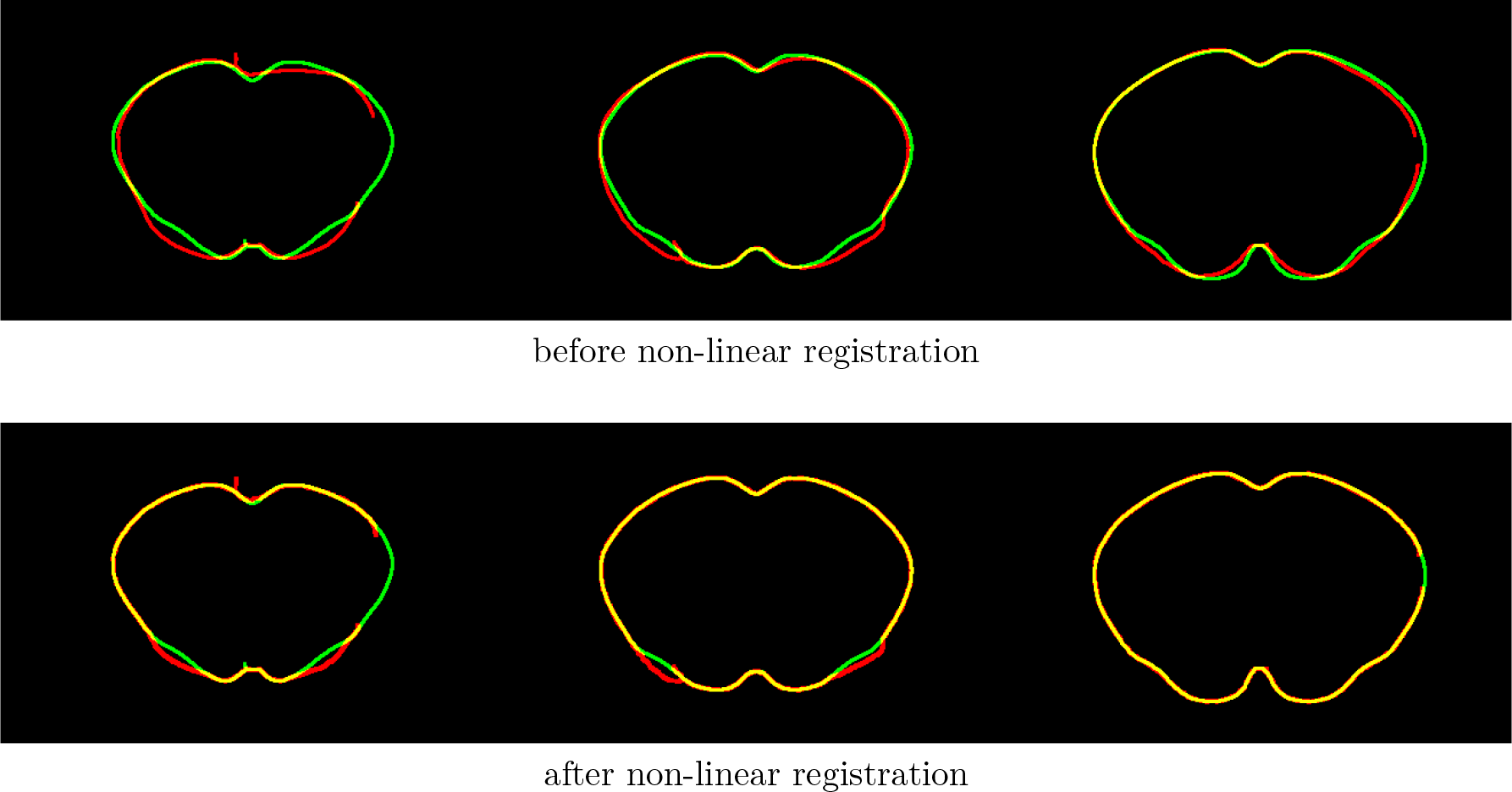
Outer contours of sample dataset planes (red) overlaid with corresponding outer contours of Atlas planes (green), before and after non-linear registration. Yellow color indicates overlap.

Having applied the transformations to the masks and quantification channels (or puncta locations) along the registration process, at last the dataset is ready for quantification.

We then read the Atlas label map, resize and crop it appropriately. If quantifying puncta, we build a volume and approximate (*x*, *y*, *z*) puncta locations by their voxel location in order to perform counting – to which we proceed. A quantification table, one line per label region, is saved as a CSV file.

## 3 Remarks

### Fast track mode

Once the pipeline is executed once, step by step, there’s the option to re-run it in *fast track* mode. This is useful, for example, if the user wants to quantify both diffuse signal and puncta: since only one type of quantification is possible at each run, the first would be done step by step, setting parameters and quantifying one type of signal, and the second in *fast track* mode, quantifying the other.

### Parallel processing

In many steps the algorithm runs tasks in parallel, which speeds up the process considerably. This requires Matlab’s Parallel Processing Toolbox. However the code should run even if the toolbox is not available – albeit using a single processor, naturally.

### Intermediate steps record

Results of intermediate steps are saved to file for recording purposes. The output names (either folders or files) always start with the name of the folder containing the dataset. Output data is saved in the same folder where the input folder is located.

### Non-linear registration

Optionally, non-linear registration can be performed on the edge maps, as in previous registration steps. The use of outer contours, as described in the text above, occurs when the variable for ‘interior contour dependence’ is active.

### Machine learning

In pre-processing, we used machine learning models to obtain masks and contours. Code and instructions for training similar models is released with the package containing code for registration, in case the user wants to adopt the same approach.

### Alternative approaches

It is a standard practice in scientific reports to only discuss the final version of the working method. For the record, however, we would like to point out some alternative, abandoned techniques. (1) We originally intended to perform 3D registration, but opted for 2D slice-by-slice registration and alignment due to its relative simplicity and the fact outputs are easier to inspect. (2) An earlier version of the pipeline used external (i.e. not built in Matlab) tools for non-linear registration; no comparison of accuracy was made, the Matlab solution being adopted for simplicity.

### Nomenclature

The toolkit is named after the card shuffling technique^4^, based on resemblance with the task it is designed to perform (half the ‘deck’ of planes is the target dataset, the other half is the Allen Atlas, and shuffling – if performed by a magician – intercalates planes according to the correct assignment). Alternative naming attempts based on acronyms failed.

## Footnotes

1 ^©^ 2019 Allen Institute for Brain Science. Allen Mouse Brain Atlas. Available from: http://mouse.brain-map.org/

2 The MathWorks, Inc., Natick, Massachusetts, United States.

3 https://www.mathworks.com/matlabcentral/fileexchange/41938-fit-2d-gaussian-with-optimization-toolbox

4 https://en.wikipedia.org/wiki/Shuffling#Riffle

## Notes

https://github.com/hms-idac/RiffleShuffle

